# Sequence-to-expression approach to identify etiological non-coding DNA variations in P53 and cMYC-driven diseases

**DOI:** 10.1101/2022.12.05.519089

**Authors:** Katherine Kin, Shounak Bhogale, Lisha Zhu, Derrick Thomas, Jessica Bertol, W. Jim Zheng, Saurabh Sinha, Walid D. Fakhouri

**Author notes:** These authors contributed equally to this work (Katherine Kin, Shounak Bhogale, Lisha Zhu).

## Abstract

Disease risk prediction based on DNA sequence and transcriptional profile can improve disease screening, prevention, and potential therapeutic approaches by revealing contributing genetic factors and altered networks. Despite identifying many disease-associated DNA variants through genome-wide association studies, distinguishing deleterious non-coding DNA variations remains poor for most common diseases. We previously reported that non-coding variations disrupting cis-overlapping motifs (CisOMs) of opposing transcription factors significantly affect enhancer activity. Analyzing publicly available ChIP-seq data for P53 and cMYC in human embryonic stem cells and mouse embryonic cells showed that ∼344-366 genomic regions are co-occupied by P53 and cMYC. We identified, on average, two CisOMs per region, suggesting that co-occupancy is evolutionarily conserved in vertebrates. Therefore, we designed *in vitro* experiments to uncover the significance of the co-occupancy and competitive binding and inhibition between P53 and cMYC on target gene expression. We found that treating U2OS cells with doxorubicin increased P53 protein level while reducing cMYC level. In contrast, no change in protein levels was observed in Raji cells. ChIP-seq analysis showed that 16-922 genomic regions were co-occupied by P53 and cMYC before and after treatment, and substitutions of cMYC signals by P53 were detected after doxorubicin treatment in U2OS. Around 187 expressed genes near co-occupied regions were altered at mRNA level according to RNA-seq data. We utilized a computational motif-matching approach to determine that changes in predicted P53 binding affinity by DNA variations in CisOMs of co-occupied elements significantly correlate with alterations in reporter gene expression. We performed a similar analysis using SNPs mapped in CisOMs for P53 and cMYC from ChIP-seq data in U2OS and Raji, and expression of target genes from the GTEx portal. We found a significant correlation between change in motif-predicted cMYC binding affinity by SNPs in CisOMs and gene expression. In conclusion, our study suggests a generally applicable approach to filter etiological non-coding variations associated with P53 and cMYC-dependent diseases.

**Author Summary:** Most DNA variants associated with common complex diseases fall outside the protein-coding regions of the genome, making them hard to detect and relate to a function. Although many computational tools are available for prioritizing functional disease risk variants outside the protein-coding regions of the genome, the precision of prediction of these tools is mostly unreliable and hence not close to cancer risk prediction. This study brings to light a novel way to improve prediction accuracy of publicly available tools by integrating the impact of cis-overlapping binding sites of opposing cancer proteins, such as P53 and cMYC, in their analysis to filter out deleterious DNA variants outside the protein-coding regions of the human genome. Using a biology-based statistical approach, DNA variants within cis-overlapping motifs impacting the binding affinity of opposing transcription factors can significantly alter the expression of target genes and regulatory networks. This study brings us closer to developing a generally applicable approach capable of filtering etiological non-coding variations in co-occupied genomic regions of P53 and cMYC family members to improve disease risk assessment.

## Introduction

Genome-wide association studies (GWAS) of different cancer types have demonstrated that around 10% of disease-associated single nucleotide polymorphisms (SNPs) are located in coding sequences, whereas 90% of the disease-associated SNPs are outside the coding regions [1-4]. Despite identifying hundreds of cancer-associated non-coding DNA variants [1, 4, 5], the ability to carry out a genetic risk assessment of diseases from these non-coding DNA variants is still lacking. By exploring potential biological mechanisms between non-coding DNA variants and disease development, we hope to address this gap, which can ultimately improve effective disease screening and prevention, and help develop novel treatment strategies.

One such mechanism of interest is competitive binding and inhibition at cis-overlapping motifs (CisOMs) because it could dictate the outcome of target gene expression. Within the non-coding genomic regions, transcription factors (TF) bind to regulatory elements with high affinity at consensus binding motifs. However, the preference for TF binding to these binding sites is influenced by several factors, including the binding affinity, the kinetics of occupancy, stoichiometry of homo-binding sites, orientation of binding sites, cooperativity, and quenching effects of neighboring TF binding sites [6-11]. In addition, the presence of cis-overlapping motifs (CisOMs) within regulatory elements suggests competitive binding and inhibition, where the binding of one factor inhibits the binding of another factor to dictate gene readout [12-14].

Competitive binding and inhibition at CisOMs have been detected in eukaryotic organisms, and previously studied in Drosophila, specifically for the transcription factors Snail and Twist [15, 16]. Notably, the mutations in CisOMs of Ebox and AP1 significantly impacted gene expression more than mutations at non-overlapping binding sites in human HCT116 cancer cells [17]. Despite these reports, the importance of CisOMs in gene regulation and the prevalence of co-occupied regions by crucial transcription factors remains to be explored at a genome-wide level in humans.

This study focuses on CisOMs involving P53 and cMYC for several reasons. First, the cellular importance of competitive inhibition on gene expression dynamics can be critical if the competition involves cis-acting transcriptional activators and repressors that are master regulators and have an antagonistic effect on the target genes. Carriers of deleterious DNA variants within CisOMs may be at high risk of developing pathological conditions like cancer due to altered expression of target genes. Second, in a previous study, potential regulatory regions with overlapping P53 and cMYC ChIP-seq signals were identified in both human cancer cells and mouse embryonic cells. A considerable number of these co-occupied regions identified in mouse embryonic cells contain multiple cis-overlapping P53 and cMYC motifs [18]. This data suggests that the expression of target genes regulated by TFs of the P53 family (P53, P63, and P73) and the basic helix-loop-helix (bHLH) family (cMYC, TWIST1, MASH1, HIF-1α) may involve competitive inhibition at co-occupied regulatory elements at the genome-wide level. Other recent publications emphasize the significance of the cis-overlapping motifs (CisOMs) for P53 and cMYC and their competitive inhibition in regulating target genes [19-21].

Multiple studies have highlighted how regulatory DNA variations within CisOMs can disrupt competitive inhibition. The presence of SNPs in CisOMs of the tumor suppressors P53 or P63 and proto-oncogene cMYC significantly altered the expression of target genes involved in signaling transduction, chromatin modifications, and DNA damage [18, 20]. Other studies showed that regulatory mutations of conserved nucleotides disrupting CisOMs strongly affect enhancer activity than SNPs that disrupt non-overlapping binding sites [12-14]. It has been shown that HIF-1α, a member of the bHLH family similar to cMYC, binds to a P53 core DNA binding motif and potentially degrades or inhibits P53 protein during hypoxic conditions, leading to a resistance to cell apoptosis [22, 23]. Another recent study reported that stem cells could be promoted for cell proliferation and differentiation in concert with only P53 and cMYC regulatory modulation in human leukemia according to proteomics, transcriptomics, and network analysis [24]. Moreover, cMYC inhibition and P53 stabilization are necessary to successfully target the proliferation of leukemic stem cells for therapeutic approaches [24]. These studies suggest a dual regulation of target genes involved in the cell cycle and differentiation by P53 and cMYC/HIF-1α [23, 24]. In addition, our previous publication provided a molecular basis for the mechanism of regulation by the opposing TFs, P53 and cMYC, which regulate many shared target genes [18]. These findings may imply that regulatory DNA variations within CisOMs can disrupt the dual regulation by competitive binding and inhibition and can potentially increase the risk for common complex diseases, for example cancer.

There are existing computational programs that can help identify functional non-coding variants. Yet, these programs have several limitations that can be improved if we uncover additional crucial features behind the mechanism of gene regulation. Two well-known computational approaches to detect non-coding functional variants include GWAVA and CADD. Both programs use machine learning that analyzes regions from previous GWAS studies to distinguish between harmful and benign variants [25, 26]. Pattern recognition and differentiation from background signals are essential methods in computational biology. Still, this approach currently does not explain the molecular mechanisms or alteration of target genes leading to diseases associated with non-coding variants. Newer approaches like FunSeq2 rely on combining different annotations like conservation of sites, loss or gain of transcription factor motifs, recurrence of variants within samples, regulatory elements to gene linkages, and the importance of affected genes within gene interacting networks to determine the effect of non-coding variants [25]. Another tool for detecting functional non-coding variants is HaploReg. HaploReg does not provide a priority score suggesting functional or benign non-coding variants. Instead, this tool provides information about the SNP itself like; its location in potential regulatory element, in the binding site, disruption of binding affinity, and location within quantitative trait loci (QTL) [27, 28]. On the other hand, FunSeq, GWAVA, and CADD computational tools provide priority scores that would prioritize functional vs. nonfunctional non-coding variants. For CADD and FunSeq, a score value larger than 20 indicates functional non-coding variations [29]. A score above 0.5 indicates functional or etiological non-coding variations in GWAVA [26].

Even with these multiple computational approaches, the sensitivity (predicted benign variants/ total number of variant effects) for analyzing and predicting functional variants varies from method to method, especially CADD, based on a recent study [29]. Current computational approaches lack the use of an expression model to predict the quantitative rather than qualitative effect of variants on gene expression leading to complex diseases, such as cancer [26]. P53 is the most mutated gene in cancer (>50% of all types of cancer), while cMYC is overexpressed in many types of cancer [24, 30]. CisOMs of P53 and cMYC, thereby are promising targets for detecting etiological variants using a computational approach that can be expanded to include P53 and cMYC family members.

We analyzed ChIP-seq data from published studies for P53 and cMYC, which showed a similar number of co-occupied regions and CisOMs between human embryonic stem cells and murine embryonic cells. To study CisOMs of P53 and cMYC in the same cells, we used U2OS and Raji cancer cells untreated and treated with doxorubicin. We used computational analysis of our ChIP-seq data to identify a substantial number of P53 and cMYC co-occupied regions before and after treatment. RNA-seq analysis showed the differently expressed genes (DEGs) after treatment with doxorubicin and we bioinformatically identified the DEGs that are close to co-occupied regions by P53 and cMYC. We further propose a biology-based sequence-to-expression statistical approach capable of assessing etiological non-coding DNA variations. We mapped putative regulatory elements with important molecular features bound by P53 and cMYC, including the number of binding sites within 100bp, CisOMs and binding affinity of each site. We also included luciferase assay data of regulatory elements that contain SNPs within CisOMs for P53 and cMYC in the statistical approach model. Therefore, this study is an important step toward developing a statistical approach or a filter to uncover etiological DNA variation for essential transcriptional factors, such as P53 and cMYC, that determine cell fate and are often dysregulated in cancer [24, 30].

## Results

### Analysis of publicly available ChIP-seq data of P53 and cMYC genomic binding in similar cell types

To identify co-occupied genomic regions containing CisOMs for P53 and cMYC, we initially used previously published ChIP-seq studies that experimentally mapped P53 and cMYC binding regions in similar cell types. We analyzed the ChIP-seq data for P53 and cMYC in human embryonic stem cells (hESC) and murine embryonic cells (mECs) (Fig 1A). We performed the analysis for 4968 P53 peaks from Akdemir, Jain [31] and 6407 cMYC peaks from ENCODE to locate co-occupied regions in human embryonic stem cells. Bioinformatics analysis showed that 366 genomic regions are co-occupied by both factors, and they contain 602 CisOMs for P53 and cMYC. According to the dbSNP database of human genetic variations obtained from UCSC and 1000 genomes, 1334 SNPs are found within the CisOM elements.

**Fig 1.**
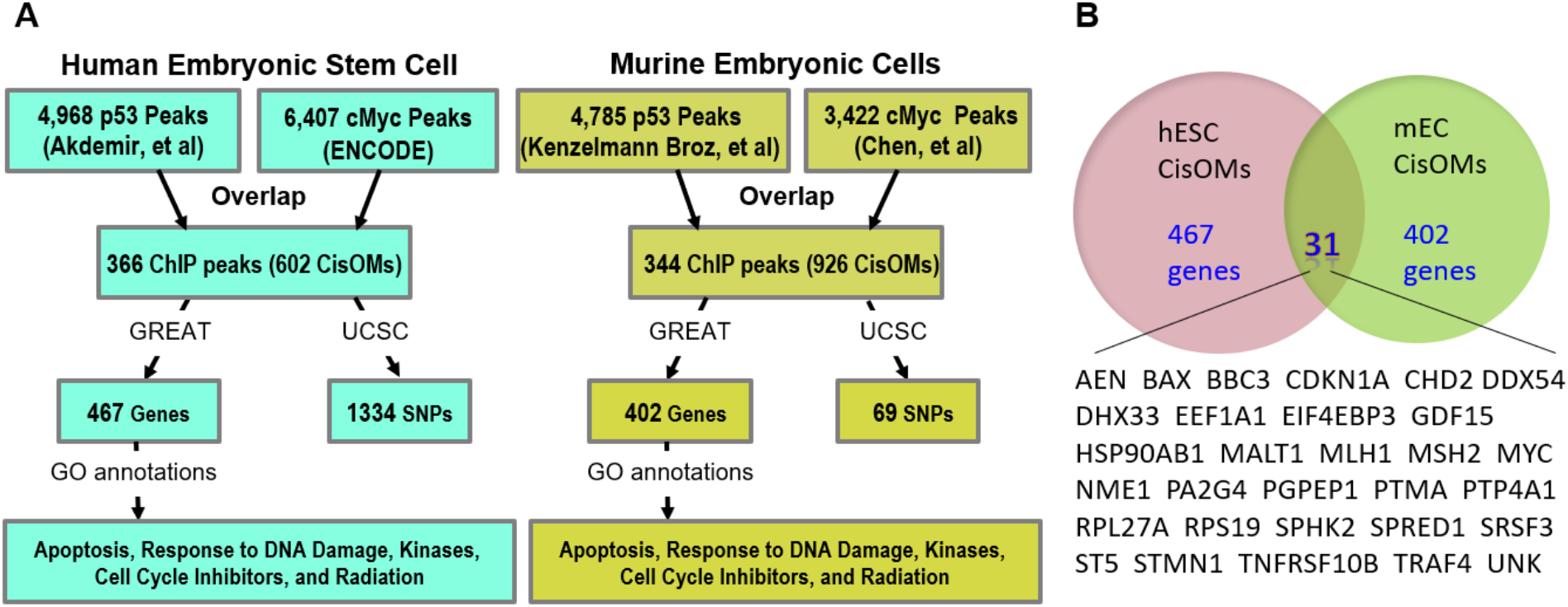
Analysis of publicly available P53 and cMYC ChIP-seq data. The flowchart illustrates how p53 and cMYC co-occupied regions from ChIP-seq data selected from studies in human embryonic stem cells and murine embryonic stem cells were used to map CisOMs of p53 and cMYC (A). CisOMs within co-occupied regions were identified using a python script. GREAT was used to reveal the potential target genes regulated by the CisOMs and USCS browser was used to identify SNPs within the CisOMs. Common Genes Regulated by COMs of P53 and cMYC in human embryonic stem cells and murine embryonic cells (B).

In murine embryonic cells, 344 overlapping peaks were identified from 4785 P53 peaks from Kenzelmann Broz, Spano Mello [32] and 3422 cMYC from Chen, Xu [33]. The co-occupied regions contain a total of 926 CisOMs for cMYC and P53, and a total of 69 SNPs within CisOMs of mouse genome. Notably, 31 genes are potentially regulated by P53 and cMYC co-occupied regions with CisOMs in both human and mouse embryonic cells (Fig 1B). According to gene ontology analysis using DAVID [34], these genes are involved in apoptosis, DNA damage repair, kinases, phosphatases, cell cycle inhibitors, and cMYC pathway.

### *In vitro* experiments for detection of P53 and cMYC in the same cancer cells

Occupancy at regulatory elements is cell-type and protein-level dependent. Therefore, we sought to determine the dynamics of the competitive binding between P53 and cMYC in two cancer cell types. We used the U2OS osteosarcoma cells and Raji Burkitt’s lymphoma cells because previous ChIP-seq data were conducted in these cell lines. Optimized experiments were conducted in U2OS and Raji cell lines to identify genomic cMYC and P53 binding signals before and after treatment with doxorubicin (Dox), a DNA-damaging drug (Fig 2A). The total protein level of cMYC and P53 were measured in cells pooled from multiple culture plates before and after treatment with Dox to assess the protein amount (Fig 2B). The level of cMYC and P53 did not change in Raji cells after Dox treatment. On the other hand, the level of P53 was remarkably increased, and that of cMYC was reduced in U2OS after Dox treatment. We then used the pooled cancer cells untreated and treated with Dox for the RNA-seq and ChIP-seq analysis.

**Fig 2.**
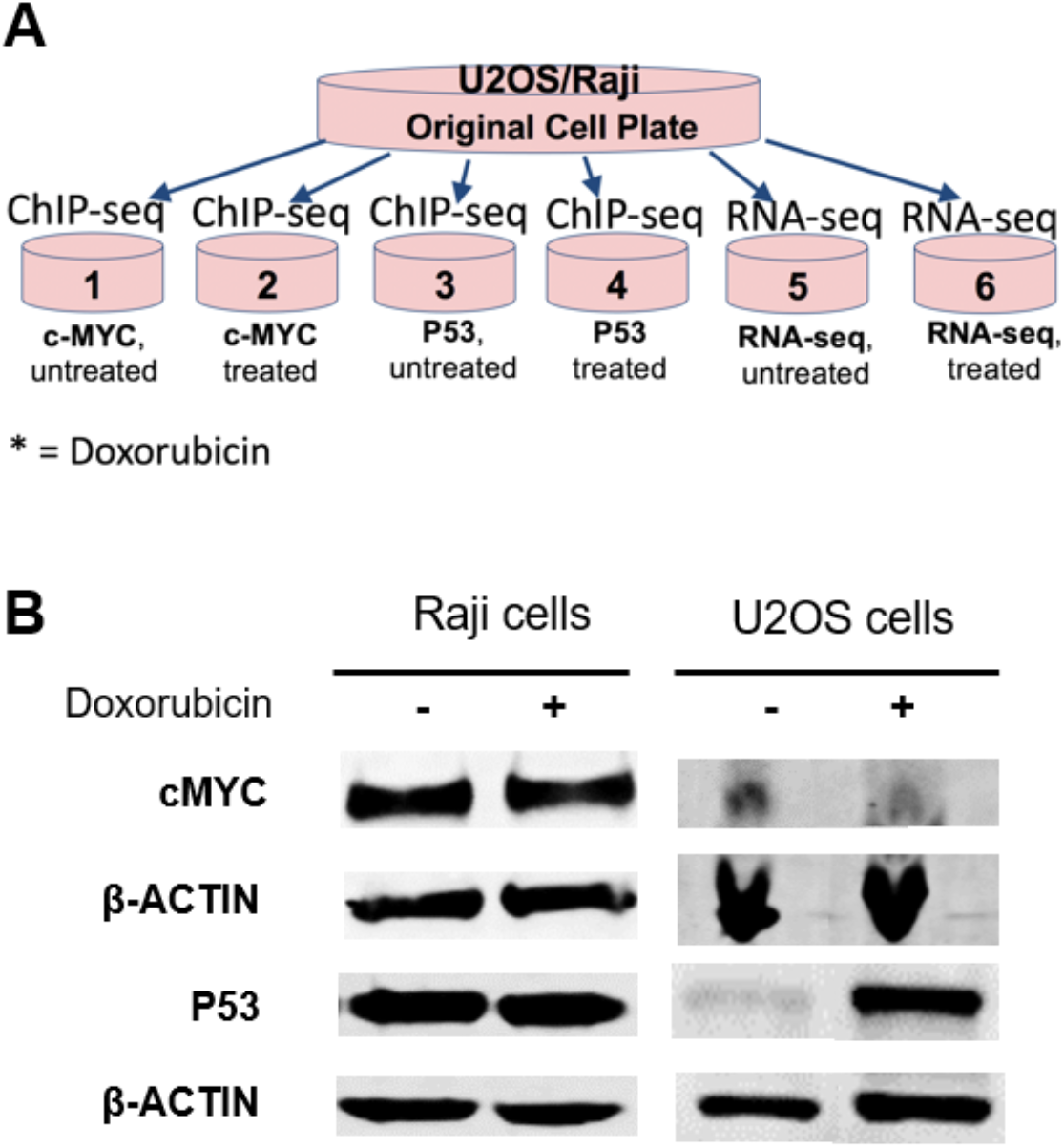
Experimental approach for computational modeling detection of P53 and cMYC *in vitro*. (A) Low passage of U2OS and Raji cells were cultured in large culture plates and the pooled cells were divided into six individual experimental Petri dishes as indicated. (B) The amount of the protein level of cMYC and P53 in U2OS and Raji cell lines was quantified after normalization to β-ACTIN before and after treatment with Doxorubicin.

### Transcriptional profile before and after treatment with doxorubicin

RNA-seq was performed to determine the differentially expressed genes (DEGs) after treatment with doxorubicin in U2OS and Raji cells. The transcriptional profile aims to link the DEGs to the co-occupied genomic regions by P53 and cMYC in U2OS and Raji cells (Fig 3A and B). We used three biological replicates for each cell line before and after treatment. The RNA-seq was conducted for 12 samples, and the transcriptional profile was generated for each sample to determine homogeneity among biological replicates (S1 Fig). Due to heterogeneity, we excluded one of the three biological replicates of U2OS-Dox treated cells. After Dox treatment, around 470 genes were upregulated and 173 genes were downregulated in U2OS cells, while 304 genes were upregulated and 599 genes were downregulated in Raji cells (Table 1). Gene ontology analysis of the DEGs in U2OS showed that these genes are involved in cell cycle, nuclear and cell division, DNA damage response, mitotic metaphase progression, and sister chromatid cohesion (Table 2). Similarly, gene ontology analysis of DEGs in Raji cells showed that these genes are involved in cell division, apoptosis, cell cycle regulation, DNA damage and inflammatory response, nuclear division, and negative regulation of transcription from RNA polymerase II promoter (Table 3).

**Table 1:** Number of upregulated and downregulated genes after treatment with doxorubicin using RNA-seq analysis

| Cell type | Upregulated Genes | Downregulated Genes |
| --- | --- | --- |
| U2OS | 470 | 173 |
| Raji | 304 | 599 |

**Table 2:** Gene ontology analysis of U2OS differently expressed genes

| Cellular Terms | Count | Percentage % | P-Value |
| --- | --- | --- | --- |
| Cell Cycle: G2/M Checkpoint | 7 | 1.33 | 1.11E-04 |
| Cell division | 31 | 5.92 | 6.31E-09 |
| DNA damage response, signal transduction by p53 mediator resulting in cell cycle arrest | 13 | 2.48 | 4.18E-08 |
| Mitotic metaphase plate congression | 10 | 1.91 | 2.65E-07 |
| Mitotic nuclear division | 22 | 4.20 | 1.50E-06 |
| Sister chromatid cohesion | 13 | 2.48 | 1.16E-05 |
| Signaling pathway | 18 | 3.44 | 2.19E-12 |
| Cell cycle | 15 | 2.86 | 8.98E-06 |

**Table 3:** Gene ontology analysis of Raji differently expressed genes

| Cellular Terms | Count | Percentage % | P-Value |
| --- | --- | --- | --- |
| Cell division | 38 | 5.35 | 8.97E-10 |
| Apoptotic process | 49 | 6.91 | 5.24E-09 |
| Regulation of cell cycle | 20 | 2.82 | 3.64E-08 |
| Cellular response to DNA damage stimulus | 26 | 3.66 | 4.12E-08 |
| Inflammatory response | 36 | 5.07 | 8.18E-08 |
| Positive regulation of apoptotic process | 30 | 4.23 | 4.14E-07 |
| Mitotic nuclear division | 26 | 3.66 | 1.21E-06 |
| Negative regulation of transcription from RNA polymerase II promoter | 50 | 7.05 | 2.78E-06 |

**Fig 3.**
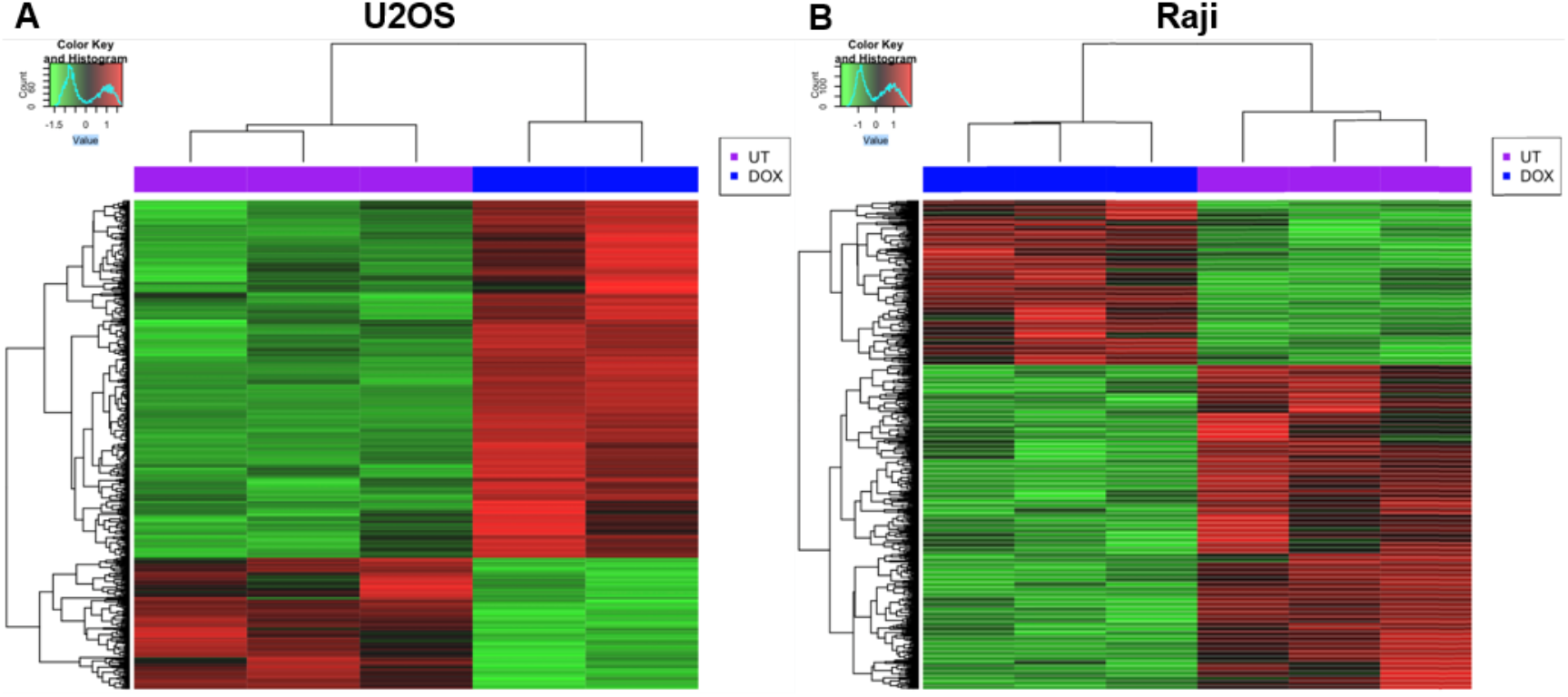
Heat map of DEGs in U2OS and Raji cells. A heat map of differentially expressed genes (DEGs) after treatment with Dox in U2OS (A) and Raji cells (B). The green color represents downregulated genes compared to control cells, while the red represents upregulated genes compared to control cells.

### ChIP-seq data analysis before and after doxorubicin treatment

ChIP-seq was performed to identify the genomic regions bound by cMYC and P53 before and after treatment with Dox in the same cancer cells. The number of ChIP peaks found within Raji and U2OS cells before and after treatment with doxorubicin is summarized in Table 4, that the number of peaks went down substantially for both TFs in Raji and went up for both TFs in U2OS. ChIP-seq analysis showed regions where additional signals of P53 were detected due to Dox treatment and regions showing a reduction of signal enrichment of cMYC after treatment which is consistent with the increased level of P53 post Dox treatment. We selected a few regions as representative of four different binding scenarios. First, in some cases, we saw an enrichment of P53 genomic binding signals due to Dox treatment in Raji cells (Fig 4A, dashed box).

**Table 4:** Number of cMYC and P53 ChIP peaks before and after treatment with doxorubicin in Raji and U2OS cell lines.

| Cell Line | Treat | cMYC Peaks<br>(*P*-value Cutoff 0.05) | P53 peaks<br>(*P*-value Cutoff 0.05) |
| --- | --- | --- | --- |
| Raji | UT | 1173 | 11316 |
| Raji | Dox | 85 | 2438 |
| U2OS | UT | 212 | 208 |
| U2OS | Dox | 320 | 805 |

**Fig 4.**
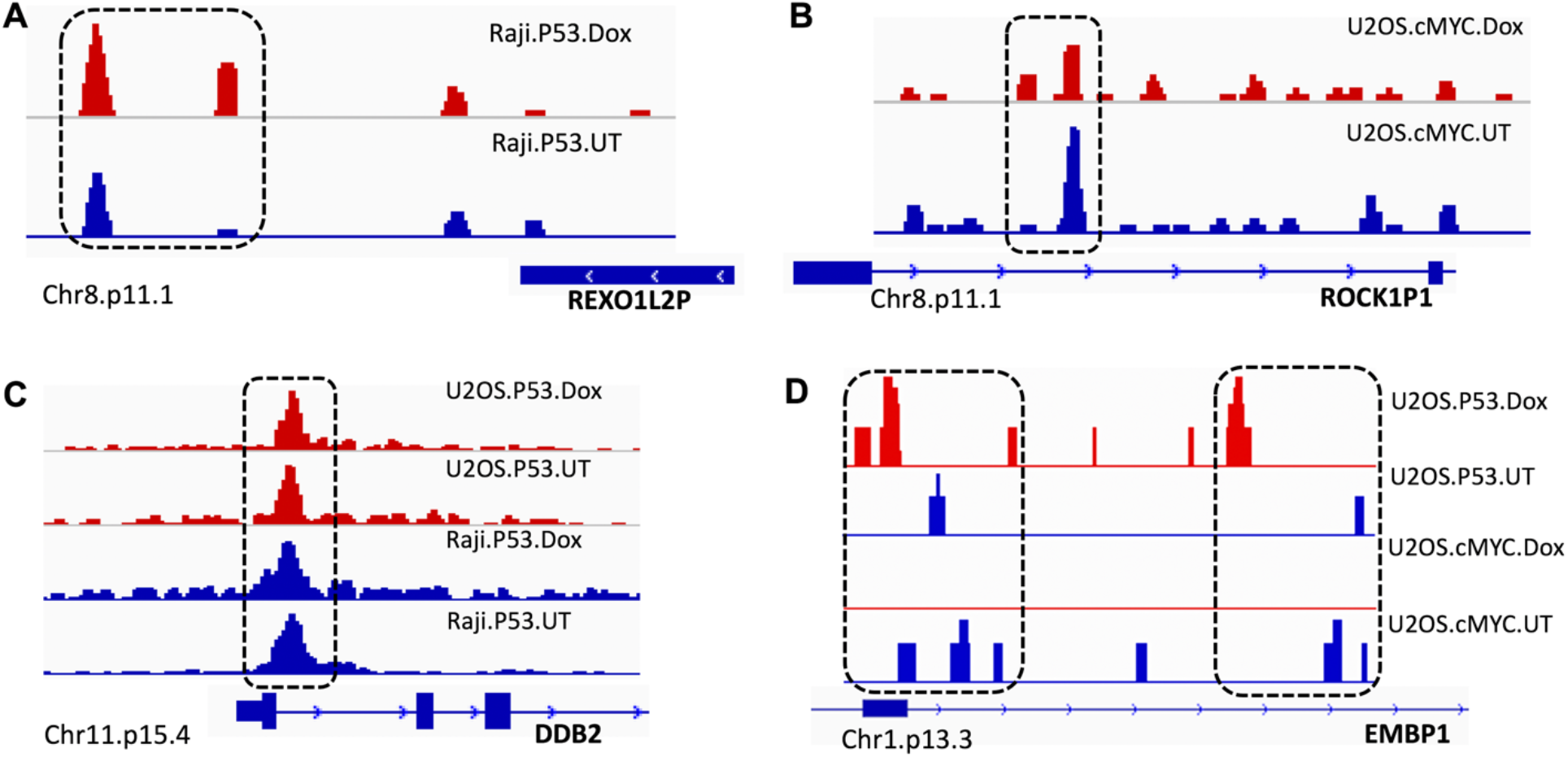
Chip-seq before and after treatment with Dox in U2OS and Raji cells. ChIP-seq was performed to identify the binding genomic regions of cMYC and P53 before and after treatment with Doxorubicin. An increased signal of p53 after doxorubicin treatment can be seen on Chr8.p11.1 in Raji cells, signals inside dotted box (A). Two increased signals and one reduced signal of cMYC can be seen on Chr8.p11.1 in U2OS cells (B). No significant changes in P53 signals can be seen on Chr11.p15.4 in U2OS and Raji cells after treatment with doxorubicin (C). However, increased signals of P53 can be seen on Chr1.p13.3 in U2OS cells and decreased signals for cMYC after Doxorubicin treatment. (D).

Second, increased cMYC signals was detected in a few regions around the ROCK1P1 intronic region in U2OS cells, however, the signal intensity was reduced (Fig 4B, dashed box). Third, there were cases where there was no change in intensity of P53 signals in U2OS cells before and after treatment with Dox, and there was a variable increase in P53 signals in Raji cells as well (Fig 4C, dashed box). Fourth, in some cases, new P53 signals and replacement of cMYC signals can be detected in U2OS cells within the same regions after treatment with Dox (Fig 4D, dashed box). However, these are not all possible scenarios of P53 and cMYC binding and enrichment. The number of co-occupied regions was bioinformatically assessed under these conditions.

### Putative DNA variations within CisOMs for P53 and cMYc generated using the PWM model

The position weight matrix (PWM) model generated all possible DNA variants within CisOMs for P53 and cMYC per potential target gene and chromosome from the Raji and U2OS experimental data. This approach aims to determine the distribution and frequency of putative DNA variations that can disrupt P53 and cMYC competitive binding per chromosome and neighboring target genes within CisOMs genome-wide (Fig 5). In untreated and treated Raji cells with Dox, we observed a relatively large number of tentative DNA variants within the CisOMs of cMYC and P53 near RUNX3, SYF2, AHDC1, FGR, SECISBP2, SEMA4D, POU2F2, CD4, LAG3, and ID2 with > 200 putative variants within CisOMs per region compared to the rest of the identified co-occupied regions (Fig 5A). In addition, we observed an enrichment in the number of tentative DNA variants within CisOMs in Chr1 in untreated Raji cells (Fig 5B). After treatment with doxorubicin, we mapped a moderate number of tentative DNA variants within the CisOMs in close proximity to ZBTB17, MTRNR2L12, TXNDC15, and in Chr1 and Chr3 in Raji compared to the other identified genomic CisOMs regions (Fig 5C and D). In U2OS cells before treatment with doxorubicin, the number of tentative variants within CisOMs is large in the PLEC gene (∼400 putative variants per region) (Fig 5E). In addition, these tentative DNA variants are enriched in Chr8 a treatment (Fig 5F). Similarly, in U2OS cells after treatment with doxorubicin, there is an enrichment in the number of tentative DNA variants (> 200 putative variants per region) in the HGH1 gene (Fig 5G). Notably, there is an increased number of variants in chr7 and chr8 (Fig 5H).

**Fig 5.**
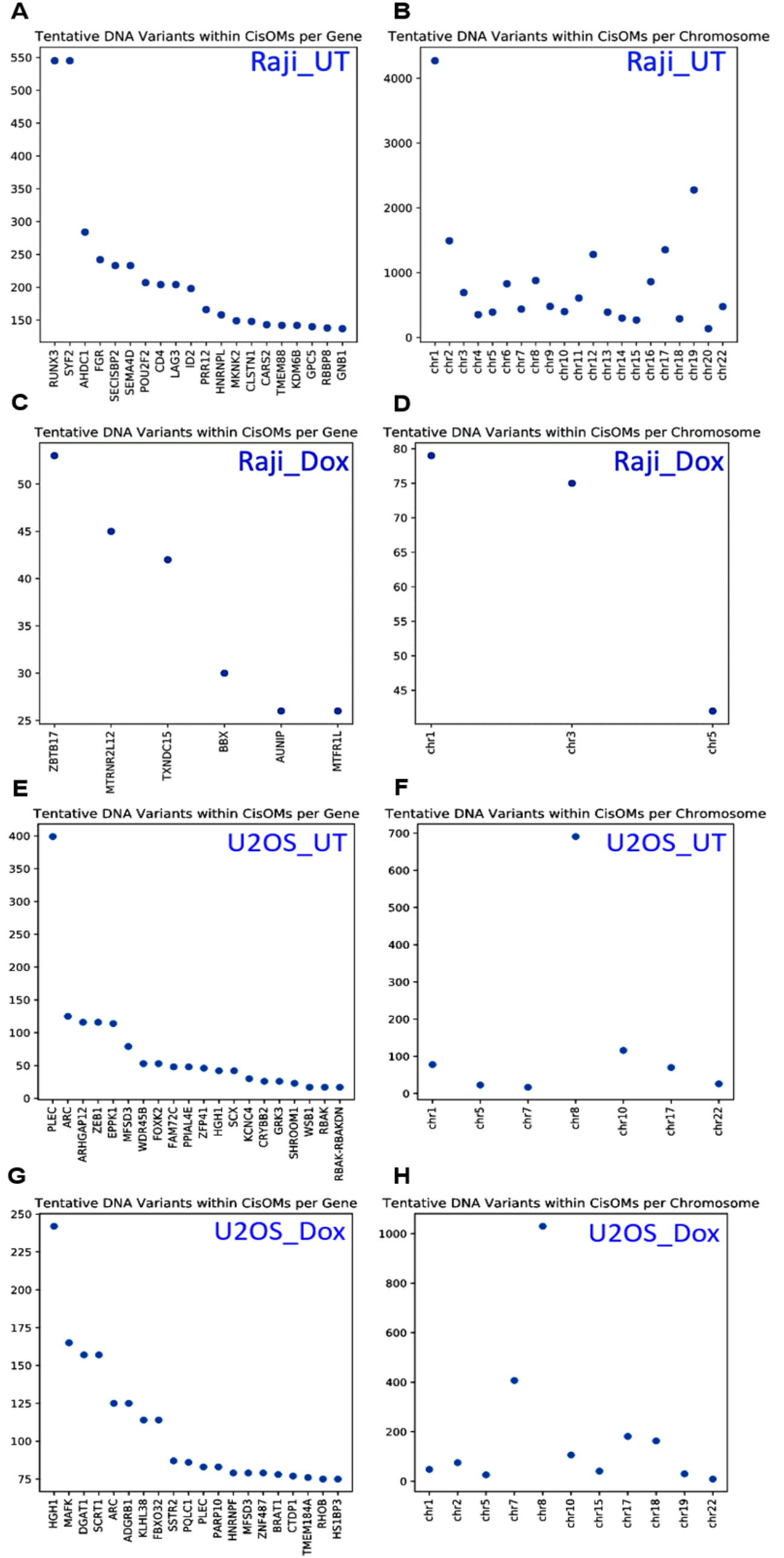
Tentative DNA variations generated using PWM model. Tentative DNA variants were mapped for CisOMs in co-occupied genomic regions by cMYC and P53 in Raji cells untreated and treated with doxorubicin. The y-axis denotes the number of DNA variants in the genes or chromosomes listed on the x-axis.

### A statistical approach to identify etiological non-coding variants in co-occupied regions by P53 and cMYC

We decided to assess 100 bp regions around the identified CisOMs as potentially minimal regulatory elements containing a SNP within CisOMs for P53 and cMYC to generate a statistical modeling approach. An illustrative diagram shows the position of the SNPs within CisOMs of co-occupied elements by P53 and cMYC (Fig. 6A).

**Fig 6.**
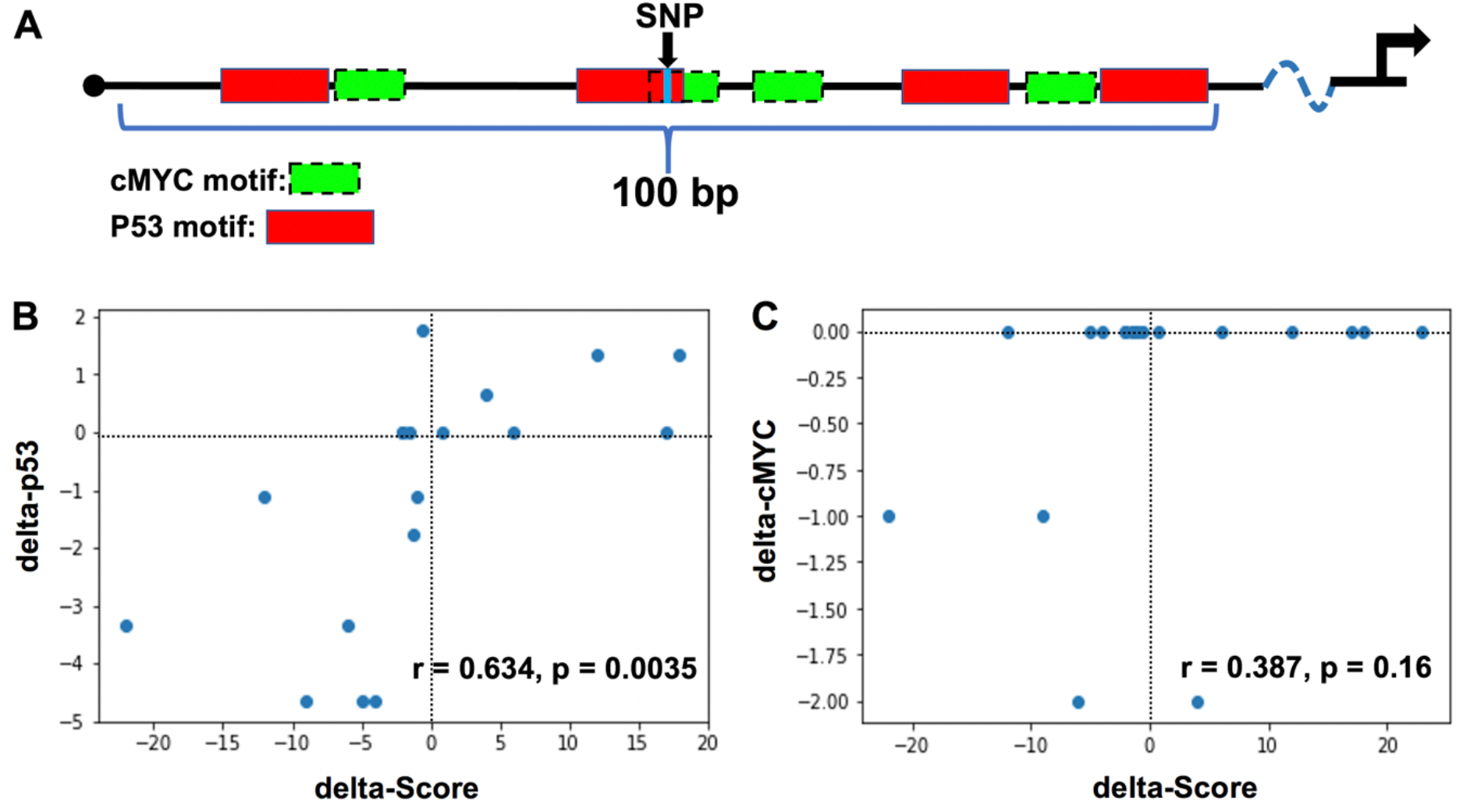
Correlation coefficient analysis for binding affinity to normalized luciferase assay score. An illustrative model shows the computational approach to generate the binding affinity to gene expression model (A). Green boxes indicate the cMYC binding site and red boxes indicate the p53 binding site. Model showing how binding strength and expression based on luciferase assay can be used to create a predictive model for p53 (B) and cMYC (C).

Luciferase assay data from previous [18, 21] and the current study was used as a readout of the P53 and cMYC co-occupied element activity. For the statistical approach, the selected SNP is located at the center of the element within the CisOMs for P53 and cMYC (Fig 6A, black arrow). For each SNP, a “delta score” was defined as the variant element-driven expression of luciferase activity minus the common/reference sequence element-driven expression of luciferase activity, thus reflecting the expression impact of the SNP. The delta-P53 scores were computationally estimated based on the change in predicted binding strength (“delta binding”) due to an SNP as the difference in motif binding score of the variant and reference alleles (Fig 6B, S2 Table). On average for the ChIP-seq data, a total of about 9.59 cMYC and 6.98 P53 binding sites in Raji cells, and 8.5 cMYC and 10.5 P53 binding sites in U2OS, were mapped within 100 bp co-occupied regions which were considered a minimal enhancer element (S1 Table). The correlation coefficient observed between altered luciferase expression (delta-score) and disruption of P53 binding affinity (delta binding) by SNPs mapped within CisOMs was significant with a p-value of 0.00355. Delta-cMYC binding strength was plotted with the delta score of the luciferase expression (Fig 6C). However, there is no noticeable pattern with cMYC data with most tested SNPs exhibiting no change in predicted binding affinity. To further expand this statistical approach to *in vivo* data, SNPs within identified CisOMs of P53 and cMYC in Raji and U2OS were used to identify likely target genes and obtain normalized gene expression values (for both alleles) from the GTEx database. Normalized gene expression values associated with 34 SNPs were obtained for common homozygous alleles and SNP heterozygous variants identified in the *in vitro* experiments (S3 Table). Expression from the homozygous alternative alleles was not used for correlation due to a lack of normalized expression values for many of the analyzed SNPs in GTEx portal. Delta expression (delta-score) and delta binding scores were calculated and used for correlation coefficient analysis. An insignificant correlation was observed between altered binding affinity of P53 and the expression of genes associated with the SNPs within CisOMs obtained from our ChIP-seq data and GTEx portal (Fig 7A, S2 Fig). However, a significant correlation between delta-cMYC and delta-score was obtained with a p-value of 0.0163 (Fig 7B). Repeating the same correlation analysis using CisOM-SNPs that have zero delta-binding scores, we also observed a significant association pattern with a p-value of 0.00515 (S2 Fig). To illustrate the impact of SNPs within CisOMs compared to non-CisOMs, normalized gene expression associated with SNPs within regulatory elements containing single (non-overlapping) binding sites of p53 and cMYC were collected for SNPs in chromosome 8 from coordinate 48007568 to 96260804. The normalized expression of 67 SNPs of the heterozygous and homozygous common alleles was obtained from the GTEx database (S4 Table). There is no significant correlation between the delta expression scores and delta binding scores for either p53 or cMYC when restricting the set of SNPs to those located within a single binding site of p53 or cMYC respectively (Fig 8A and B, S3 Fig).

**Fig 7.**
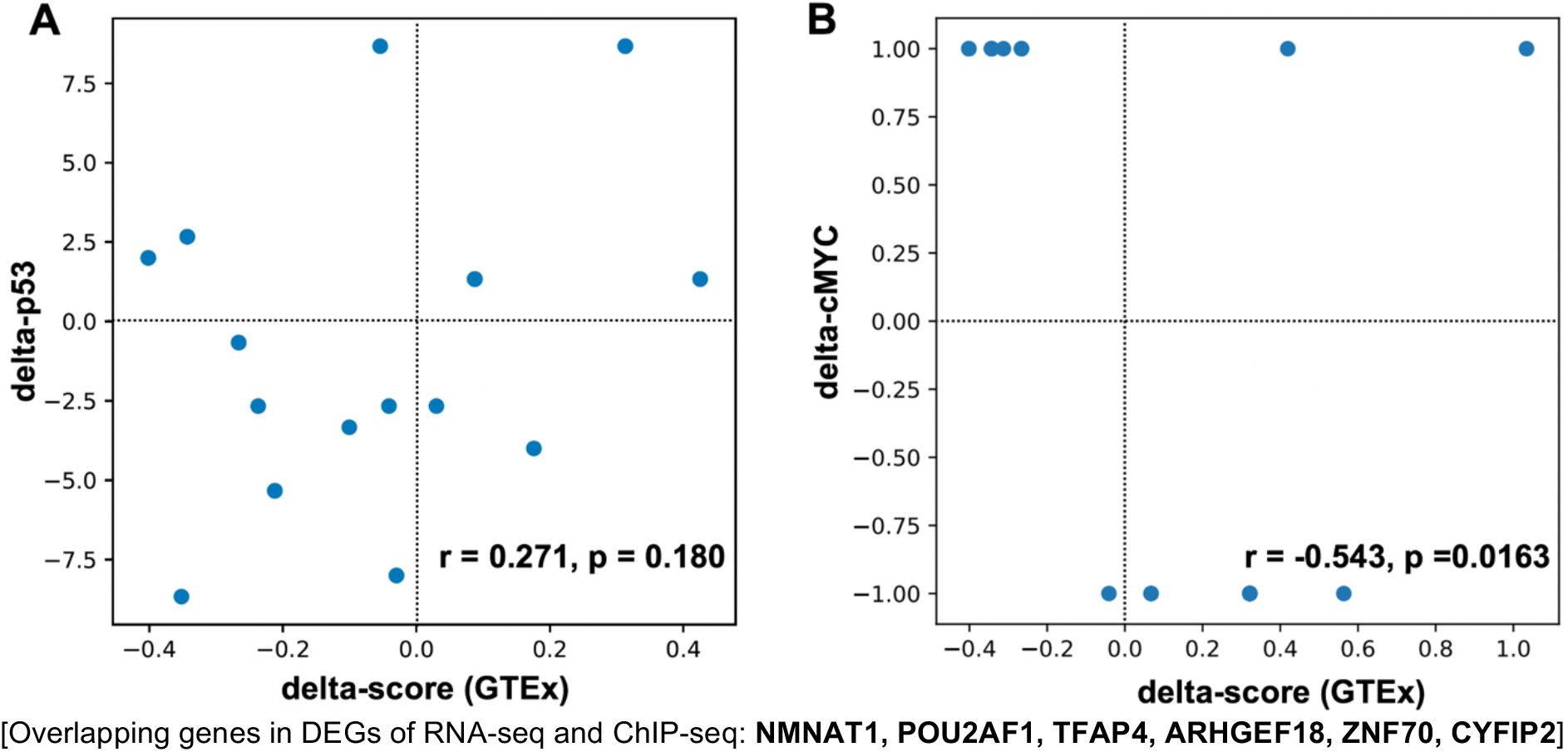
Correlation coefficient analysis for binding affinity to normalized expression score in CisOMs. The statistical approach generated using expression level data from GTEx database shows how binding strength and normalized expression can be used to create a predictive model for p53 (A) and cMYC (B) using SNPs within CisOMs of p53 and cMYC.

**Fig 8.**
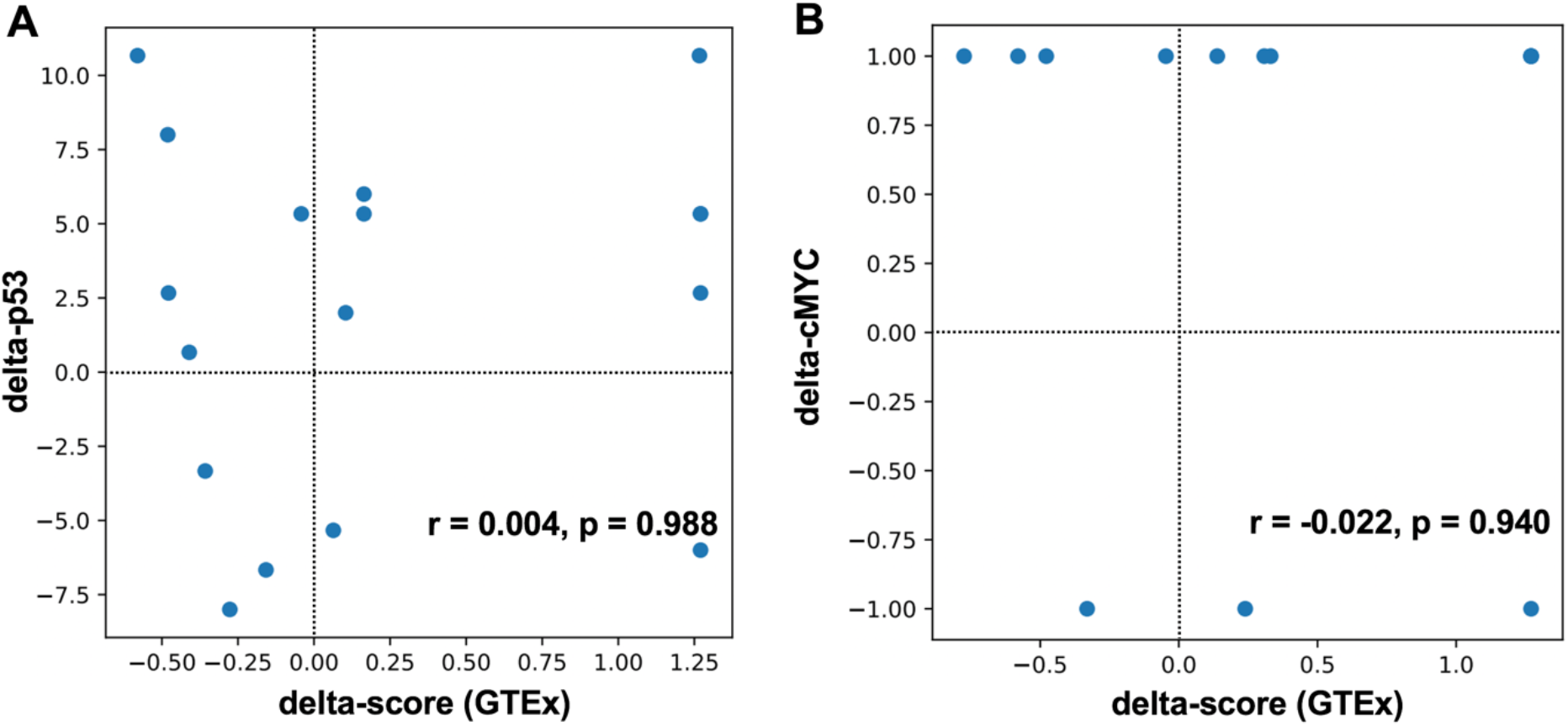
Correlation coefficient analysis for binding affinity to normalized expression score in single binding sites. The statistical approach was generated using expression level data from GTEx database showing how binding strength and normalized expression can be used to create a predictive model for p53 (A) and cMYC (B) using SNPs within single binding sites of cMYC and p53.

## Discussion

There are several factors for the poor performance of predictive computational models of functional non-coding DNA variants, such as the inadequate understanding of the molecular feature of gene regulation, the inaccurate estimation of the DNA variant impact on target gene expression, the significance of DNA sequence conservation in vertebrates on the functional regulatory element, and enrichment of transcription factor binding sites. In cancer diseases, risk prediction based on non-coding variants remains challenging due to the diversity of the non-coding regions among individuals, the inability to distinguish driver and passenger mutations, and the current lack of understanding of the underlying mechanism associated with functional non-coding variants [35]. Hence, proving an underlying biological mechanism for disease development, such as the functional non-coding SNPs.

To help tackle these limitations, this study used a well-controlled and optimized *in vitro* experiment to validate the importance of P53 and cMYC competitive binding and inhibition within CisOM elements. We hypothesize that P53 and cMYC competitive binding and inhibition at co-occupied regions are crucial for regulating expression level of target genes, and that SNPs located within CisOMs have more substantial deleterious effects on target genes than DNA variations in single binding sites. We investigated two crucial transcription factors crucial for cell cycle and are involved in cancer initiation, progression, and dissemination. We determined how the competitive binding of these factors can significantly impact the expression of associated genes.

Therefore, the study aimed to test the enrichment of functional/etiological SNPs within CisOMs of P53 and cMYC compared to DNA variations in non-CisOMs and single occupied regions.

Previous studies reported that HIF-1α (bHLH family member similar to cMYC) binds to P53 DNA binding motifs and that HIF-1α directly stabilizes P53 protein during hypoxic conditions, while HIF-2α suppresses it [22, 23, 36]. Another study showed that simultaneously targeting cMYC and P53 modulation can cure chronic myeloid leukemia, where cMYC inhibition, coupled with P53 stabilization, targets leukemic stem cells for elimination [24]. These studies suggest a strong interaction between P53 and cMYC to differentially regulate genes and the importance of balancing between P53 and cMYC levels to maintain cell homeostasis. Meanwhile, the findings of this study provide a molecular mechanism for how these factors with opposing cellular functions regulate target genes by co-occupied elements and propose an innovative approach to identify and predict pathological regulatory DNA variations within CisOMs in healthy and affected states.

We used ChIP-seq data from previous studies to identify the co-occupied genomic regions by P53 and cMYC in similar cell types, human embryonic stem cells (hESC) and murine embryonic cells (MEC). ChIP-seq data analysis revealed that the number of P53 and cMYC overlapping peaks identified in human embryonic stem cells and mouse embryonic cells are very similar (366 vs. 344). In addition, each co-occupied element has on average two CisOMs in both hESC and MEC. This suggests that the competitive binding feature between P53 and cMYC to regulate target gene expression is conserved in other mammalian species. A total of 31 genes were found in common between the CisOMs of hESC and MEC, where most of these genes are involved in different types of cancer. For example, mutations within the DNA mismatch repair genes, MLH1 and MSH2, can compromise the repair of mutations leading to accumulation of passenger mutations. In contrast, mutations in the MYC gene can lead to cell cycle disruption and genomic alterations [24, 37]. Several of the identified putative target genes of co-occupied regions containing CisOMs are known target genes of P53 and cMYC that are crucial for fundamental cellular functions [38-40]. Furthermore, gene ontology analysis suggested that the competitive inhibition at CisOMs between P53 and cMYC is involved in controlling apoptosis, cell cycle, response to DNA damage, and radiation in humans and mice.

To validate these findings in the same cells, the genomic binding of P53 and cMYC was studied in two cancer cell lines, Raji and U2OS, expressing P53 and cMYC. We used two cell lines, since the occupancy of the transcription factors differs among cell types. Cancer cells that express mutant P53 protein in Raji did not respond to stimulation by DNA damaging drugs because the mutant P53 protein is expressed at high levels at all stages of cell proliferation in Burkitt’s lymphoma [41]. This can be observed in our western immunoblot data where there was no difference in P53 or cMYC expression in Raji cells after treatment with doxorubicin. In contrast, we noticed a significant increase in P53 expression and a slight decrease in cMYC expression after treatment with doxorubicin in U2OS cells. Analysis of differentially expressed genes before and after treatment with doxorubicin using RNA-seq analysis showed that around 643 genes in U2OS cells and 903 genes in Raji cells were differentially expressed. These genes regulate signaling pathways, cell cycle, chromatin remodeling, and DNA damage response.

Our study mapped co-occupied regions by P53 and cMYC containing CisOMs of their motifs. It analyzed the impact of many DNA variants within the CisOMs to determine if they strongly affect target gene expression. The ChIP-seq data for the two cancer cell lines before and after treatment with doxorubicin showed four scenarios of genomic binding by cMYC and P53. First, there can be a reduction in the height of cMYC signal peak after treatment. Second, it can be substituted with other neighboring signals leading to potentially altered gene expression, third, there can be no change.

Finally, an interesting finding observed by this study is that cMYC signals can be replaced with a P53 signal, or new P53 signals can be detected after treatment with doxorubicin in regions near cMYC signals. This is consistent with a previous finding where the reduction of *cMYC* expression induced by P53 binding to a downstream enhancer element of *cMYC* was important in DNA damage response [42]. Direct suppression of *cMYC* by P53 might explain the reduction of cMYC binding signal intensities in some co-occupied genomic elements. This may also explain the replacement of cMYC signal with P53 in genomic regions after Dox treatment [42].

In Raji cells, putative DNA variants generated using PWM showed that we mapped an enrichment in the number of putative DNA variants within CisOMs of P53 and cMYC in Chr1 in untreated cells, and Chr 1 and Chr 3 in treated cells with Dox. We predict an enrichment in the number of tentative variants within RUNX3, a tumor suppressor, and SYF2, a cell cycle regulator, as targets for both P53 and cMYC, where the two genes play an important role in cancer progression [43, 44]. In U2OS cells, there was a notable enrichment in the number of tentative DNA variants in chromosome 8 for P53 and cMYC regions before and after treatment with Dox which is interesting since cMYC gene is in chromosome 8. There was also an enrichment within the PLEC gene, a member of the laminin-binding integrin family and is important for cancer invasion and metastasis [45]. Many of the other identified genes as putative targets of co-occupied CisOM regions are known target genes of P53 and cMYC. These genes are correlated with P53 mutations, and are involved in different types of cancer, and cellular functions [46-50].

Finally, to generate a sequence-to-expression approach to detect functional SNPs, we performed a correlation analysis using luciferase normalized activity levels and motif-based predicted binding affinity of P53 and cMYC with or without SNPs in CisOMs of elements ranging from 606-1349bp. The data analysis showed a significant correlation between the delta luciferase activity scores and the delta binding affinity scores of P53 (p-value =0.0046). This indicates that DNA variants (SNPs) that alter the binding affinity of P53 within CisOMs compared to the common sequence result in a measurable impact on altered target gene expression. In contrast, there was no observed correlation between the change in luciferase activity score (“delta-expression”) and change in predicted cMYC binding affinity scores (“delta-cMYC binding”) because only a few variants in our dataset disrupted cMYC binding affinity. To validate these findings *in vivo*, we mapped all SNPs within CisOMs identified in our ChIP-seq data in an unbiased approach and identified 34 SNPs with putative target gene expression from GTEx portal database. Using human expression values from multiple tissues from GTEx database, an insignificant correlation (p-value = 0.180) was observed between delta-p53 binding scores and delta-expression scores for the SNPs identified within CisOMs of co-occupied regions. On the other hand, a significant correlation (p-value = 0.0163) was detected between delta-cMYC binding scores and delta-expression scores for SNPs in CisOMs. Although our analysis was performed with SNPs within P53/cMYC cis-overlapping motifs, our findings should encourage future examination with detailed characterization of the impact of SNPs within CisOMs of other members of P53 and cMYC family members on target gene expression compared to SNPs in single binding sites. This study emphasizes the need to develop a refined computational model tailored to include the significance of CisOMs on regulatory element activity and target gene expression, which could be used as a filter to map deleterious non-coding DNA variations out of a large number of disease-associated SNPs. Expansion of our approach would include the P53 and cMYC family members, such as P63, P73, TWIST1, HAND2, MAX, MyoD, HIF1, and MASH1, which emphasize the significance of DNA variations within CisOMs.

## Materials and Methods

### Detection and analysis of CisOMs

ChIP peaks for P53 and cMYC binding regions were obtained from a previous study [31] and ENCODE database for human embryonic stem cells. ChIP peaks for murine embryonic cells were also obtained from previous studies [32, 33]. We repeated the process for CisOMs analysis, gene ontology analysis, and SNP analysis as previously described [18]. Gene annotations in GREAT and DNA variant analysis by UCSC were also performed as previously described [18]. SNPs identified in hESC were from all SNPs 150 track on UCSC, GRCh38. SNPs identified in mESC are from SNPs 138 track, mm9.

### Cell culture for U2OS and Raji cells

CCL86 Raji and HTB96 U2OS cells were cultured for ChIP-seq analysis and RNA-seq analysis before and after treatment with doxorubicin. We used 150 ng/ml concentration of doxorubicin for U2OS cells and 200 ng/ml for Raji cells. The multiple culture plates were pooled and then divided into six separate Petri dishes for cMYC, P53, and RNA-seq analysis with or without treatment with doxorubicin.

### Protein analysis by western blot

Immunoblotting was performed on total protein extracts from untreated and treated Raji and U2OS cells to detect the protein levels of P53 and cMYC. We used mouse anti-P53 (sc-126) and mouse anti-cMYC (sc-40) for the immunoblot. Odyssey Li-Cor system was utilized to visualize the protein bands and intensity, and beta-actin was used as a loading control.

### RNA-Seq of Raji and U2OS transcriptional profile

RNA-Seq analysis was conducted on untreated and treated Raji and U2OS cells with doxorubicin to identify differentially expressed genes after treatment. The total RNA samples extracted from cancer cells were submitted to Genewiz company, which performed an independent test on RNA quality and concentration. Illumina HiSeq2500 platform in a 2×100bp paired-end configuration was used to obtain 35 million reads on average for each library sample, and the analysis of the DEGs was performed using the Bioconductor edgeR package. Normalized read count by the total read count per sample was converted to log counts per million (log CPM) metrics, and used for clustering and generating a heat map using the clustered image maps (CIM) function in the mixOmics R package.

### Bioinformatic analysis of RNA-Seq data

The raw reads obtained from paired-end RNA Sequencing were mapped to human reference genome hg38 using STAR [51] with default parameters to obtain count values. The annotation gtf file is gencode v28 from GENCODE [52]. The edgeR [53] was used to obtain the DEGs between untreated and treated Raji and U2OS cells with an FDR cutoff of 0.05 and a fold change cutoff of 1.5. Bioinformatic analysis was performed on the DEGs in Raji and U2OS identified by RNA-Seq analysis to determine the potential molecular function and cellular impact of these DEGs. The online tool Database for Annotation, Visualization, and Integrated Discovery (DAVID) was used to determine the gene ontology of DEGs.

### ChIP-Seq of Raji and U2OS genomic binding profile

ChIP-Seq analysis was conducted on untreated and treated Raji and U2OS cells with doxorubicin to identify genomic binding after treatment. The sonicated and cross-linked DNA samples extracted from cancer cells were submitted to Genewiz company, who performed an independent test on DNA quality and concentration. Illumina HiSeq2500 platform in a 2×100bp paired-end configuration was used to obtain 15 million reads on average for each library sample. Illumina adapters were first trimmed by Trimmomatic (v0.39) [54] and then reads were aligned to hg38 with bowtie2 [55] using default parameters. The low-quality reads (MAPQ < 10, PCR duplicates) were excluded from the further analysis. Peaks were called for each sample using MACS2 [56] with following parameters “-q 0.05 -B-nomodel--extsize 200”.

### DNA variants generated using the PWM model

We mapped P53 and cMYC binding sites in co-occupied genomic regions in Raji and U2OS cells. To generate all possible putative DNA variations at CisOMs, we used position weight matrix (PWM) of P53 and cMYC binding motifs. The overlapping ChIP peak sequences were scanned using GEMSTAT’s sequence annotator tool [57] for identifying the binding sites of p53 and cMyc. Each site is assigned a likelihood ratio (LR) score based on the PWM of the respective TF [58]. Since p53 binds as a tetramer, we used half-site PWM and allowed a variable spacer length of 2-15 base pairs between the two half-sites. Thus, two half-sites of p53 will be considered as a site only if the distance between them is less than 15 base pairs. The LR score for this combined site was calculated by taking the sum of the LR score of the individual half-sites. A “CisOM (Cis Overlapping Motif)” is a region in the overlapping peaks of p53 and Myc where a p53 full site and a cMyc site overlap with each other. The CisOM is defined as the smallest possible region which covers both these binding sites. Variant sequences were generated by mutating the base at every position in the CisOM one at a time and all three possible mutations were considered. The mutation’s effect on the binding of the two TFs was obtained by recalculating the LR score of the variant sequence.

### Site-directed mutagenesis and luciferase assay in cell culture

We used HEK293 cells for plasmid transfection in a 96-well plate with a glass bottom that contains DMEM, 10% FBS, and no antibiotics medium at 37°C as previously described [18]. The HEK293 cells were transfected 2 h after plating using lipofectamine 2000 (Life Technology, CA) with pGL3-basic-Luc and pGL3-enh-Luc as negative and positive controls, respectively, and with the co-occupied regulatory regions fused upstream of the luciferase gene. The pGL3-SV40-Renilla plasmid served as an internal control for transfection efficiency. Site-directed mutagenesis was used to introduce the identified SNPs within CisOMs as previously described using PCR and DpnI digestion enzyme [19].

### Predicted-Motif Binding Affinity of Co-occupied Elements to Expression

The overlapping ChIP peak coordinates were determined within 100bp from the center of each SNP in each direction. The delta scores for the gene expression were generated by subtracting the luciferase scores of the common allele from the alternative allele. Similarly, delta binding affinity was generated by subtracting the binding score of the common from the alternative allele of all P53 or cMYC binding sites within the 100bp of the regulatory element. The delta binding scores were obtained as described previously using the computational motif-matching approach [58] for the two transcription factors, P53 and cMYC. However, the range of the delta binding scores is different for P53 than cMYC for better display reasons. ChIP-seq data from untreated and treated Raji and U2OS cells were used to obtain SNP IDs within p53 and cMYC binding sites within CisOMs with normalized expression in the GTEx portal in other cells. The SNP IDs were filtered out to only include SNPs within the CisOMs of p53 and cMYC. SNP IDs of SNPs within CisOMs were used to collect normalized expression data from the GTEx database. Around 2000 SNP IDs were manually entered in the GTEx portal, and only 34 SNPs had normalized expression data in the GTEx portal. The tissues used in GTEx portal are typically from whole blood, skin, adipose, lungs, and others totaling up to 50 different tissue types. The tissues are obtained from healthy individuals between the ages of 20-70. Most donors are white, but the newer update includes African American individuals. Two-thirds of the sample were males. The samples were obtained post-mortem with common causes of mortality. Individuals with metastatic cancer or who had received chemotherapy were not eligible for the study.

To compare the impact of SNPs within CisOMs to non-CisOMs, the normalized expression for SNPs within single binding sites of p53 or cMYC was collected for SNPs in chromosome 8:48007568-96260804 due to a large number of SNPs. The median values of the normalized expression of genes associated with the SNPs were collected from the common homozygous, heterozygous, and alternative homozygous alleles of the analyzed SNPs. In addition, the normalized expressions were mainly obtained from the whole human blood, skin, lungs, and breast tissues. Delta-scores and delta-p53 and cMYC scores were calculated, and correlation coefficient analysis was performed using Spearman’s correlation method. A p-value less than 0.05 was considered statistically significant.

## Acknowledgements

We sincerely thank Drs. Amina Qutub and Bryan Long for their excellent contribution and mentoring of Dr. Katherine Kin during the initial stages of the computational modeling and mathematical concepts. We are grateful to Jamie Jiang, who optimized the cell line culture and genomic processing for ChIP-seq analysis. This study was funded by an R15 GM122030/NIGMS to WDF. LZ and WJZ were supported by CPRIT RP170668 and NIH 1 UL1 TR003167 01. The work was partly supported by NIH grant R35GM131819A and funds from Wallace H. Coulter Distinguished Faculty Chair in Biomedical Engineering to SS.

## Supporting Information

**S1 Table.** Number of P53 and cMYC binding sites within 100bp of co-occupied regions in Raji and U2Os.

| Cells | Treatments | n. elements | n. cMYC | n.P53 |
| --- | --- | --- | --- | --- |
| Raji | UT | 142 | 9.59 | 6.98 |
| U2OS | UT | 4 | 6.0 | 9.0 |
| U2OS | DOX | 18 | 11.11 | 12.45 |

**S1 Fig.**
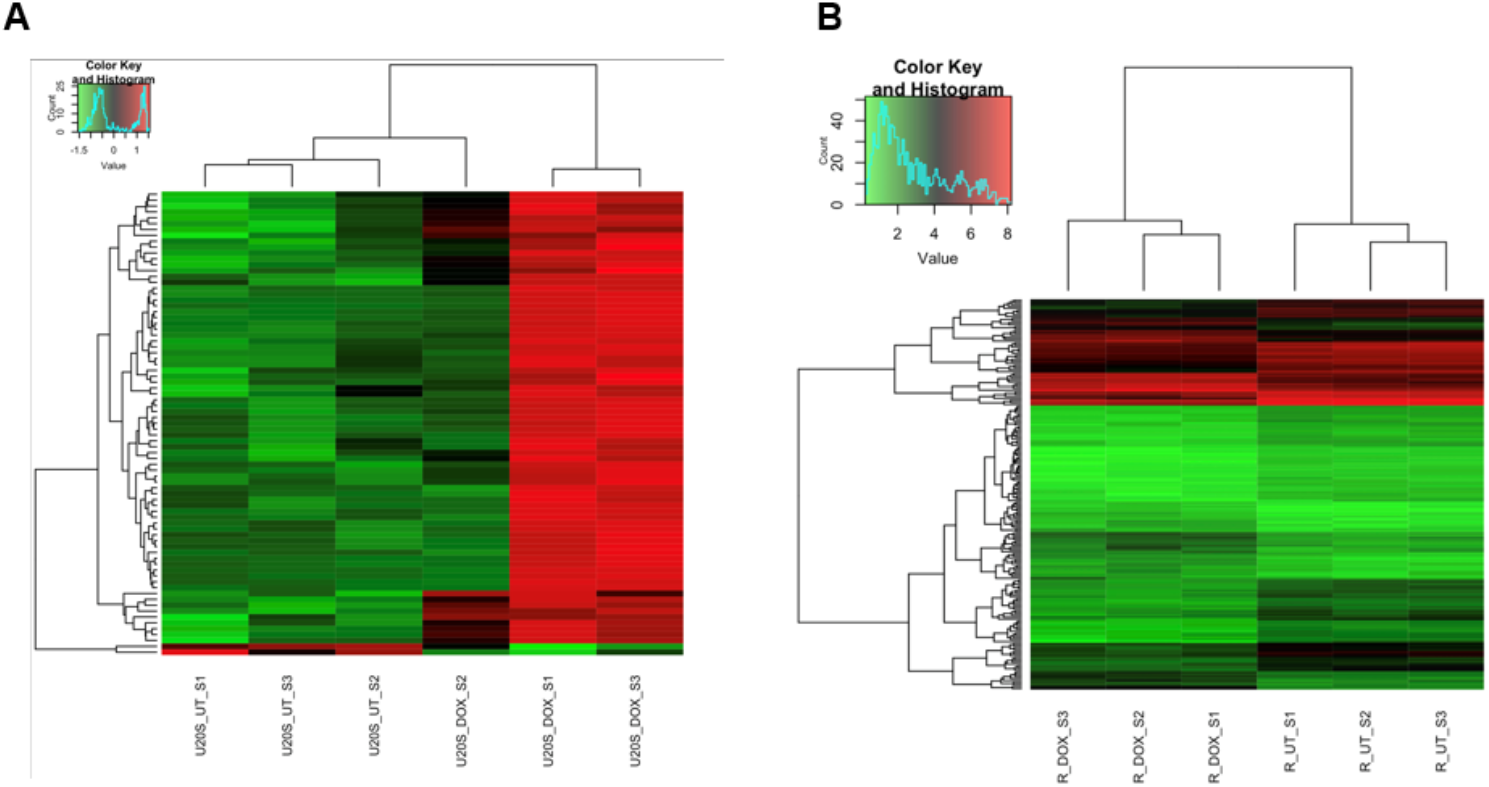
Heat map of DEGs in U2OS and Raji cells. Heat map of differentially expressed genes after treatment with Dox in U2OS (A) and Raji cells (B).

**S2 Fig.**
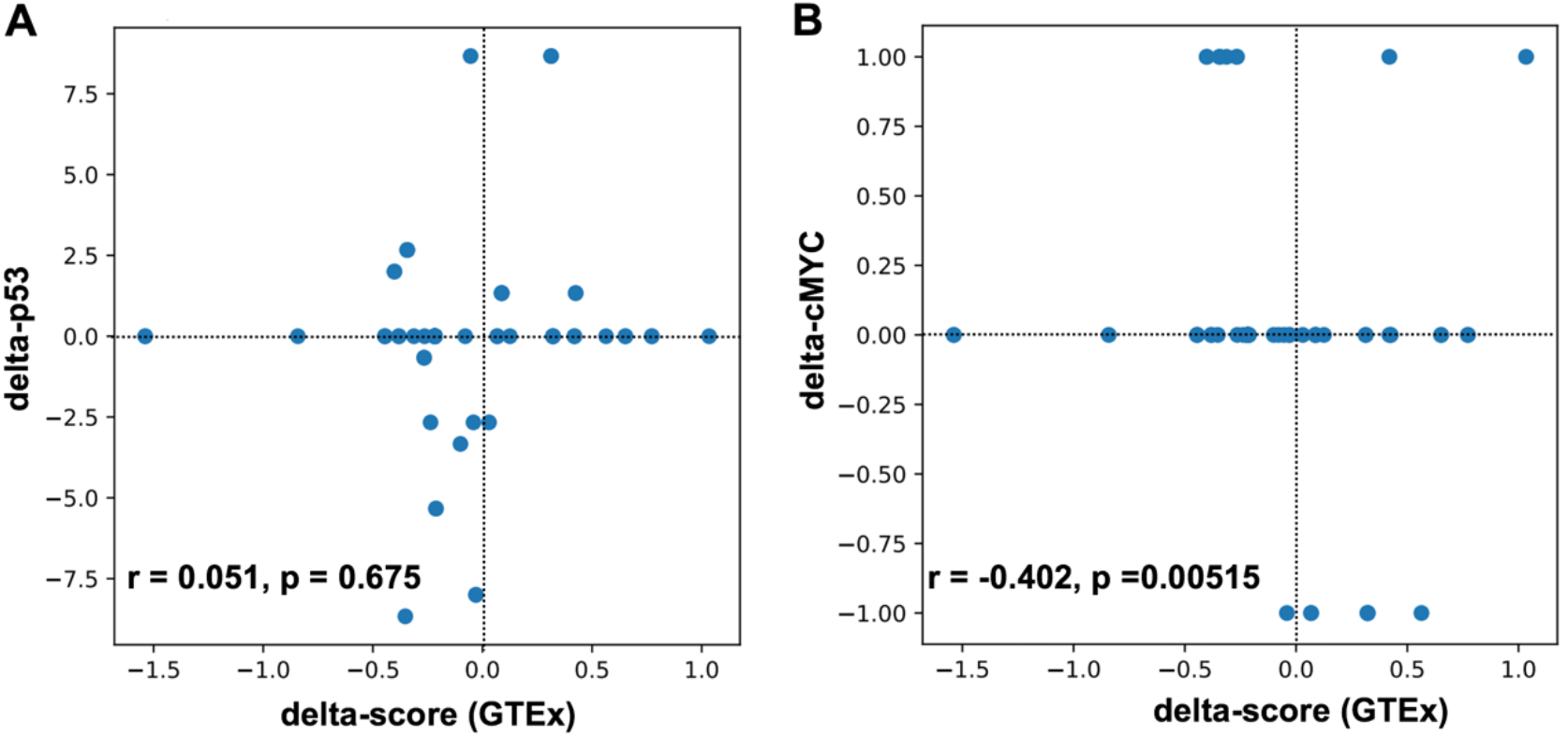
Correlation coefficient analysis for binding affinity to normalized expression score in CisOMs. Statistical approach generated using normalized expression data from GTEx database and binding strength for p53 (A) and cMYC (B) before excluding data points with no change in binding affinity.

**S3 Fig.**
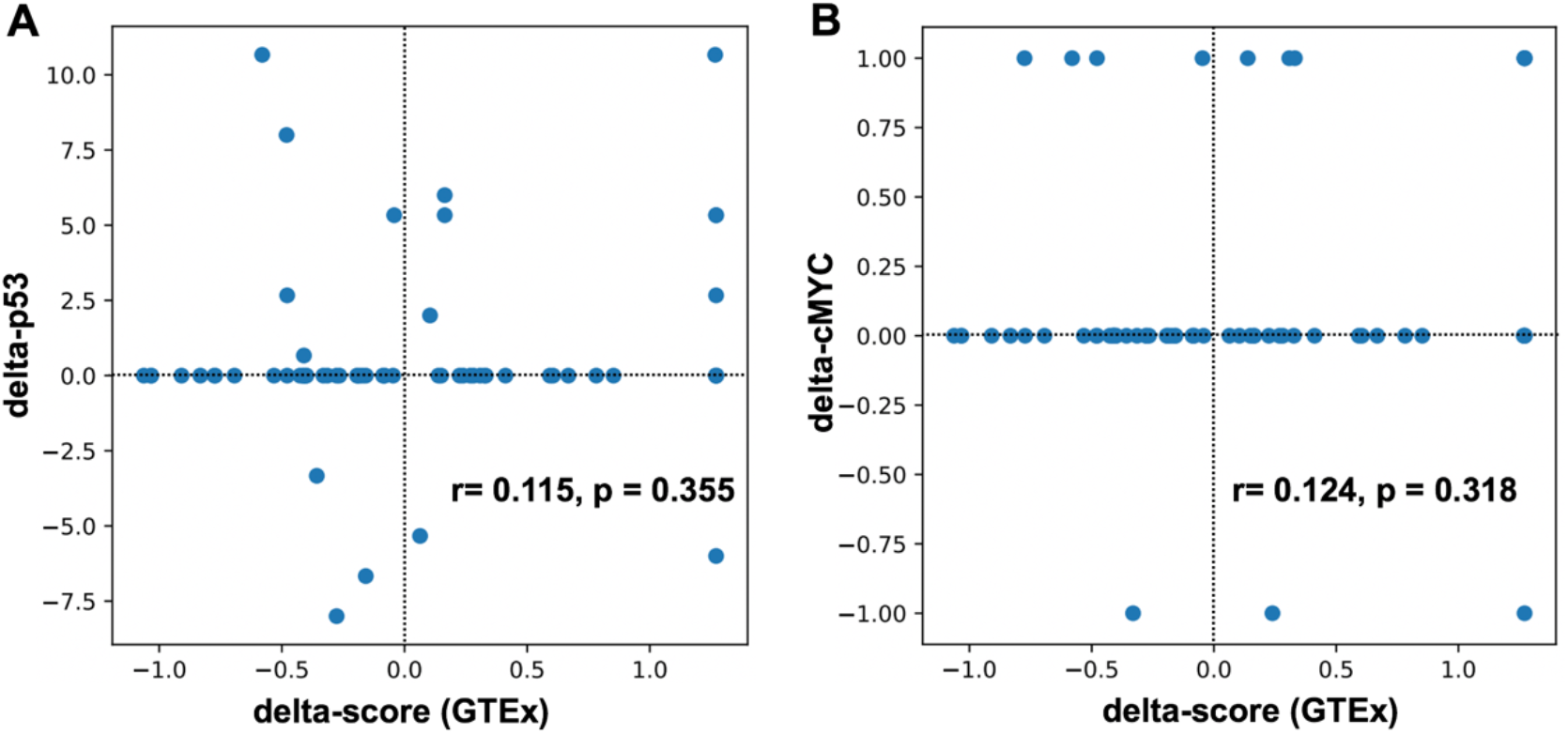
Correlation coefficient analysis for binding affinity to normalized expression score in single binding sites. The model generated using expression level data from GTEx database showing how binding strength and normalized expression can be used to create a predictive p53 (A) and cMYC (B) before excluding data points with no change in binding affinity.

